# Multiscale Rugosity Assessment of Coral Reefs Using Underwater Photogrammetry

**DOI:** 10.1101/2025.01.17.633548

**Authors:** Erick Barrera-Falcón, Rodolfo Rioja-Nieto, Roberto C. Hernández-Landa

## Abstract

Understanding the structural complexity of coral reefs is essential for assessing their condition, biodiversity, and resilience. Traditional methods commonly use a rugosity index, based on the chain method, that overlooks the underlaying structure of coral reefs. However, digital underwater photogrammetry allows to build models of coral structures which can then be used to decompose reef topography across multiple layers. This study introduces a wavelet-based method for the multiscale analysis of reef rugosity, considering reef’s surface and underlying characteristics. Data were collected from six reefs within the Cozumel Reefs National Marine Park (CRNP) at depths ranging from 6 to 14 m. High-resolution Digital Elevation Models (DEMs) and orthomosaics were constructed using digital underwater photogrammetry (UWP). The elevation profiles extracted from the DEMs were analyzed using a Maximum Overlap Discrete Wavelet Transform (MODWT), with a Daubechies mother wavelet to decompose the reef topography into local rugosity (related to live coral coverage) and underlying rugosity (related to the historical context of the formation of the reef matrix). The wavelet-based method effectively decomposed the DEMs into components representing rugosity at different scales, with the reconstructed DEMs being statistically equivalent to the data source (p> 0.05). Underlying reef characteristics contributed the most to the rugosity estimates. Significant differences in rugosity were observed between reefs (p< 0.05), where interpretations differed based on the contribution of surface and underlying characteristics. For the CRNP, Agariciid and branching corals are the primary drivers of surface rugosity (p < 0.05), rather than the mound & boulder, and meandroid corals. Our results highlight that traditional methods to estimate rugosity can underestimate the importance of local rugosity in maintaining rugosity over time.

## Introduction

The growth and spatial distribution of scleractinian corals shape coral reef seascapes [1, 2]. Geologically, coral reef development depends on a delicate balance between accretion and erosion, which is influenced by both physical and biological factors [3, 4]. This dynamic interplay regulates the growth potential of a reef through the net carbonate accretion or erosion of its calcareous matrix[5].

Assessing the structural complexity of coral reefs is crucial for understanding their ecosystem services and processes [1, 6]. Structural complexity refers to the physical architecture of a reef, including the organisms that colonize the calcareous matrix and add dimensionality [1, 7]. The structural complexity is vital for reef communities and is positively correlated with fundamental biological processes such as feeding relationships, reproduction, and competition [8, 9].

Initial attempts to measure reef structural complexity date back to the 1970s [10], when the technique of laying transects with a chain along a single linear profile over a substrate was introduced. This approach led to the development of the rugosity index as a proxy for estimating reef structural complexity [11], which is now commonly referred to as the chain method [12]. Since then, rugosity has become the most widely accepted metric for representing the structural complexity of coral reefs [13, 14]. The chain method quantifies the ratio of the distance a chain travels when laid over the substrate to the linear distance between its ends [12]. On flat surfaces, this ratio is equal to one, with values greater than one indicating increased structural complexity [11].

Alternative approaches have been suggested for measuring structural complexity, which incorporate a third dimension [15-17]. These techniques are costly, logistically challenging, and require sophisticated equipment, advanced programming knowledge, and powerful computing resources, making their widespread implementation difficult [18]. However, improvements in underwater techniques, particularly data acquisition and advancements in image analysis, have facilitated the development of methods and metrics that integrate third dimensions into measurements [14]. Close-range photogrammetry was introduced in coral reef studies by Bythell et al. (2001) to obtain structural complexity metrics for coral colonies with simple morphology. Since then, many studies have incorporated photogrammetry, hereafter referred as underwater digital photogrammetry (UWP), to analyze the structural complexity of coral reefs [19-23]. UWP allows for a more precise assessment of individual colonies in large areas [17, 24], making it a new standard in coral reef studies [25].

Despite advancements in UWP, the integration of multiscale analysis in coral reef spatial assessments remains insufficiently explored. Existing methods neither adequately capture the spatial variability of reef complexity nor effectively decompose topographic signatures to reveal the underlying structural features across different scales. Assessments of coral reef rugosity at various spatial scales were conducted by Ferrari, R. et al. (2016), Harris et al. (2023), and Ribas-Deulofeu et al. (2021). However, the spatial decomposition of topographic signatures remains unexplored. Therefore, it is important to develop a robust and scalable technique that can provide a better understanding of the complex topography of reefs and their spatial heterogeneity.

This study proposes applying a wavelet filter to high-resolution coral reef elevation profiles obtained with UWP, to analyze coral reef topography and characterize the reef structural complexity. By conducting a spatial decomposition of topographic signatures, we facilitated a comprehensive multiscale analysis [28-31]. Specifically, this study aims to (1) use a wavelet filter on high-resolution coral reef elevation profiles of six shallow reef systems of Cozumel Island and (2) conduct a spatial decomposition of topographic signatures to enable multiscale analysis.

## Materials and methods

### Study area

The Cozumel Reefs National Park is in the municipality of Cozumel, 16.5 km off the coast of the state of Quintana Roo (Fig 1). Cozumel is the second most populated island in Mexico, with 84519, inhabitants, as recorded in INEGI (2020). Its main economic activity is tourism, which is associated with SCUBA diving and cruise ships [33]. Since the 1970s, the population density and urban development along the coast have increased considerably leading to significant changes in land use and modifications to wetland ecosystems along the western margin [34].

**Fig 1.**
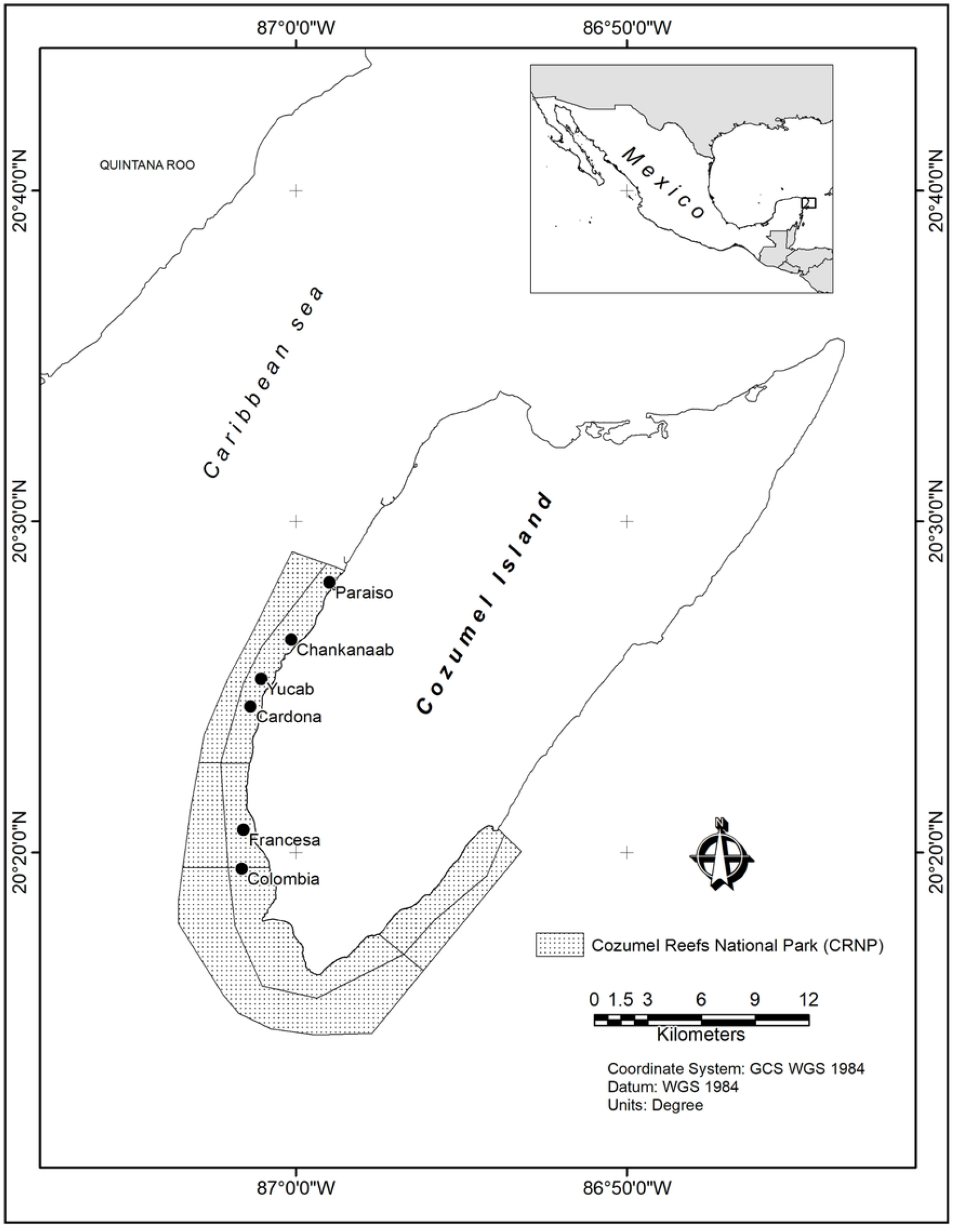
Study Area, black dots indicate the locations of the studied reefs within the Cozumel Reefs National Park. Fig 1. **In situ rugosity by reef using the chain method**. The ordering of reefs follows a north-to-south direction. The black lines represent the median values of rugosity, highlighting variation among reefs.

Marine habitats in the CRNP include a mix of marginal reefs, patch reefs, and mixed corals over hard calcareous substrates, along with algal meadows, seagrass beds, and mangrove areas [35]. The shallow sublittoral slope tends to be narrow and descends gradually from the coast, with the most developed reefs found along the edge of the southwestern insular platform [35].

### Data collection

Data were collected from six reefs within the CRNP at depths ranging from 6 to 14 m (Fig 1). Three plots measuring 5 × 30 m were established at each reef, covering a total area of 450 m^2^ per reef. The plots were delineated with markers made of 60 × 60 cm polyvinyl chloride (PVC) quadrats, placed at each corner and at the midpoint of each plot. Divers swam (along and across the plots), at c.a. 5m/s maintaining 2 m from the surface, capturing photographs with Canon G12 cameras, ensuring a high degree of overlap between adjacent images to facilitate accurate photogrammetric reconstruction [36]. Depth was recorded at the central point of each marker to provide precise elevation references.

### Photogrammetric processing, colony identification and digitization

The images from each plot were processed to construct digital elevation models (DEMs) and orthomosaics using Agisoft Metashape software v1.5. The main workflow involved aligning the images, identifying the control points (60 × 60 cm PVC markers), and applying scale corrections using known dimensions of the markers. Re-optimization of the alignment was performed to correct horizontal discrepancies, and the Z-axis was fixed by incorporating the depth data recorded from the field markers, ensuring accurate vertical scaling [36]. A dense point cloud was generated using the corrected tie points, which was then used to extract the DEMs and build an orthomosaic for each plot. The DEMs were exported in raster format by assigning an arbitrary projected coordinate system to facilitate the spatial analysis.

The orthomosaics were analyze in ArcMap v. 10.8 where coral colonies were identified, digitized, coded, and grouped according to the Atlantic and Gulf Rapid Reef Assessment (AGRRA) protocols [37]. Species identification was performed visually using the Reef Coral Identification Guide [38], Coralpedia (https://coralpedia.bio.warwick.ac.uk/), and AGRRA identification guides [37], considering colonies ≥ 5 cm, which is the minimum size that can be identified in the orthomosaics [36].

### Assessment of Rugosity

*In situ* rugosity was assessed in each plot using the chain method [39], which involves laying a chain along the contour of the substrate over a 10-meter transect and calculating the ratio of the chain length to the linear distance.

Digital rugosity was assessed in the laboratory using algorithms written in MATLAB v2020 (S1 File.zip). The height values of each pixel row or profile in the DEM were treated as time-series data and a Maximum Overlap Discrete Wavelet Transform (MODWT) was applied [40, 41]. The Daubechies wavelet was used as the mother wavelet with four levels of decomposition [42], allowing for multiscale analysis of surface features.

To evaluate the decomposition of the DEM, the original model was compared with the sum of the topographic signatures using the Mann-Whitney U test (p = 0.05). The sum of the first four decomposition levels was considered as inherent surface variations (e.g. such as those determined by coral colonies), representing small-scale features, hereafter defined as digital local rugosity (DRL). The approximation at level five was defined as the underlying structure, representing larger-scale relief without inherent variations, defined as the digital global rugosity (DGR). This approach is analogous to creating a digital terrain model that excludes the surface roughness, focusing solely on a broader underlying matrix.

For each profile, digital rugosity was estimated using the roughness parameter Ra, as defined by Raja, Muralikrishnan and Fu (2002). Ra, represents the arithmetic mean of the absolute deviations of the profile height from the mean line over a specified evaluation length (5 cm), calculated using equation 1.

To standardize the digitization process and extract precise measurements of Ra, an evaluation length of five centimeters was used to assess both DRL and DGR profiles. This length corresponds to the minimum diameter of digitized coral colonies.

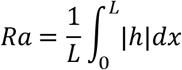

Where:

h = absolute height of surface profile at each sampling pixel.

L = evaluation length.

The results were stored in the same spatial position as the corresponding pixel, resulting in a raster that displayed spatial variations in rugosity.

### DLR and DGR based on coral cover

The vector files of the digitized coral colonies were rasterized, with each pixel tagged with a species code to enable spatial filtering. Rasterization considers the pixel size of the DRL and DGR matching the pixel size of the DEMs. These datasets were organized into a three-dimensional matrix. The matrices representing DLR and DGR occupied the first and second dimensions, respectively, whereas the raster of coral cover was placed in the third dimension.

To analyze the data, and decrease the processing time, we considered of 380 m^2^ per reef, as this is representative of the coral community of the shallow reefs in the study area [36]. Spatial overlap was performed by filtering the DLR and DGR matrix values according to site and coral groups. Since the data were not normally distributed, comparisons of digital rugosity among reefs and the contributions of coral shape groups were conducted using the non-parametric Kruskal-Wallis test (H test, p = 0.05). When the test indicated significant differences, pairwise comparisons between sites were performed using Dunn’s post-hoc test with the Bonferroni correction [43].

## Results

The reconstruction using the wavelet approach and the decomposition in DLR and DGR, is show in in Fig 2. In all cases the reconstructed DEMs were equivalent to the original DEM (p > 0.05; S2 File), indicating that wavelet filtering effectively decomposes the DEM for rugosity analysis in both the DLR and DGR. Furthermore, the DGR is higher than the DLR.

**Fig 2.**
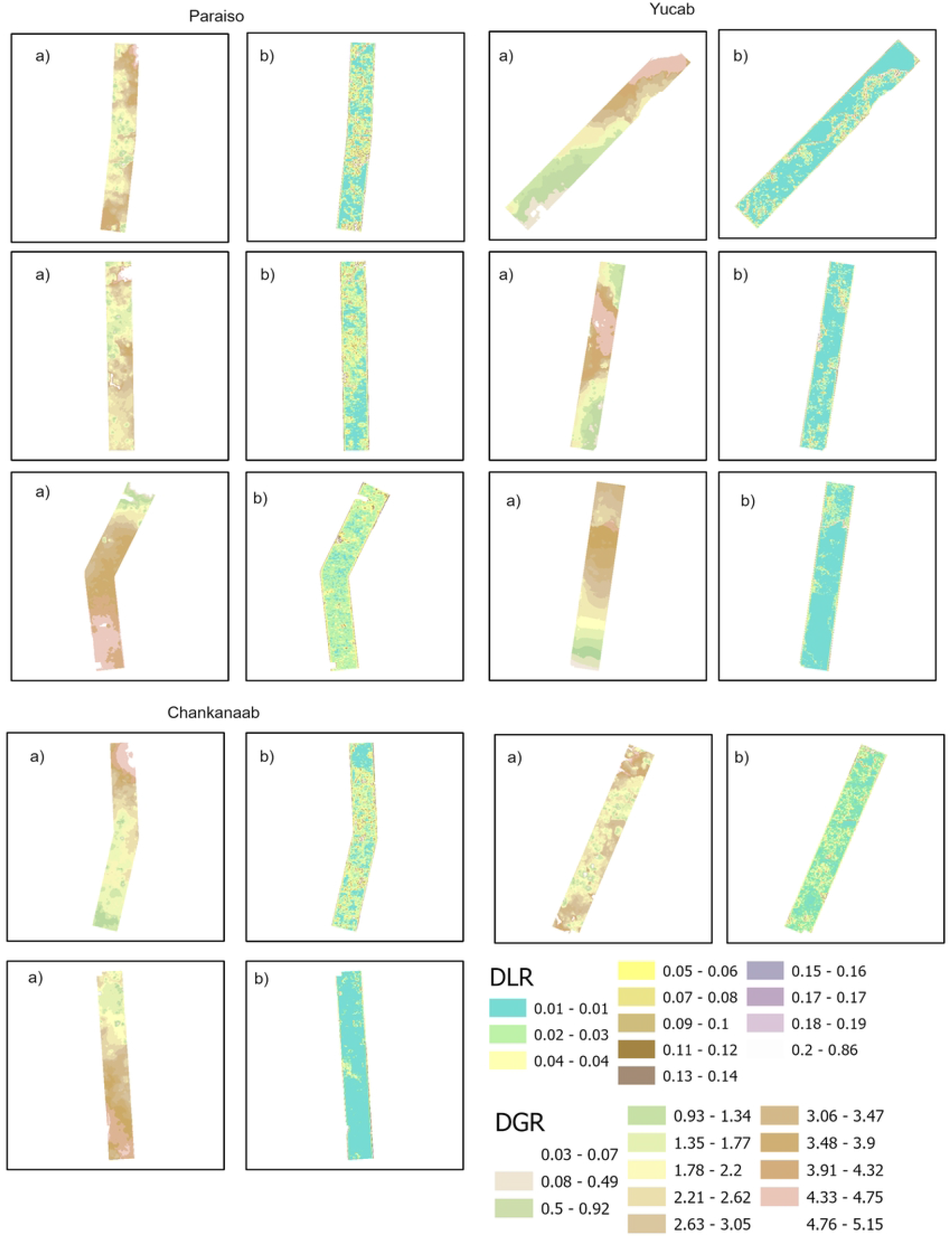
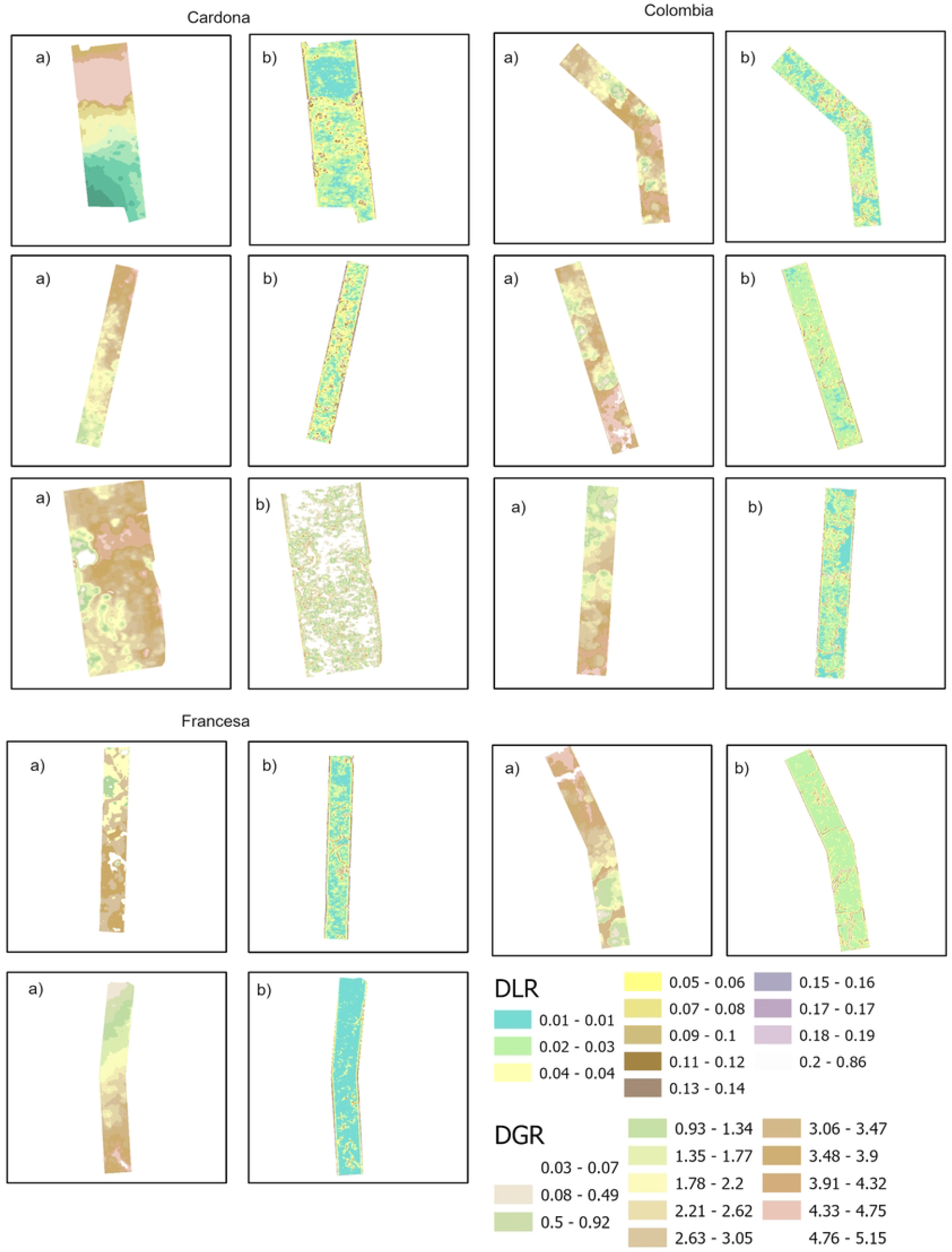
Total rugosity (DGR + DLR) per reef. The ordering of reefs follows a north-to-south direction. This integrates both the DGR representing the structural component shaped by long-term historical processes and DLR which reflects the contribution of living coral and benthic organisms.

Fig 2a. **Digital roughness values of Paraiso, Yucab and Chankanaab reefs**. (a) DGR: highlighting the historical contributions to reef structure, (b) DLR: emphasizing the biological contributions from live coral communities.

Fig 2b. **Digital roughness values Cardona, Colombia and Francesa reefs.** (a)DGR: highlighting the historical contributions to reef structure, (b) DLR: emphasizing the biological contributions from live coral communities.

*In situ*, the rugosity showed that Francesa and Colombia have higher values than Yucab, Chankanaab, Cardona and Paraiso, where the later showed the lowest values (Fig 3).

**Fig 3.**
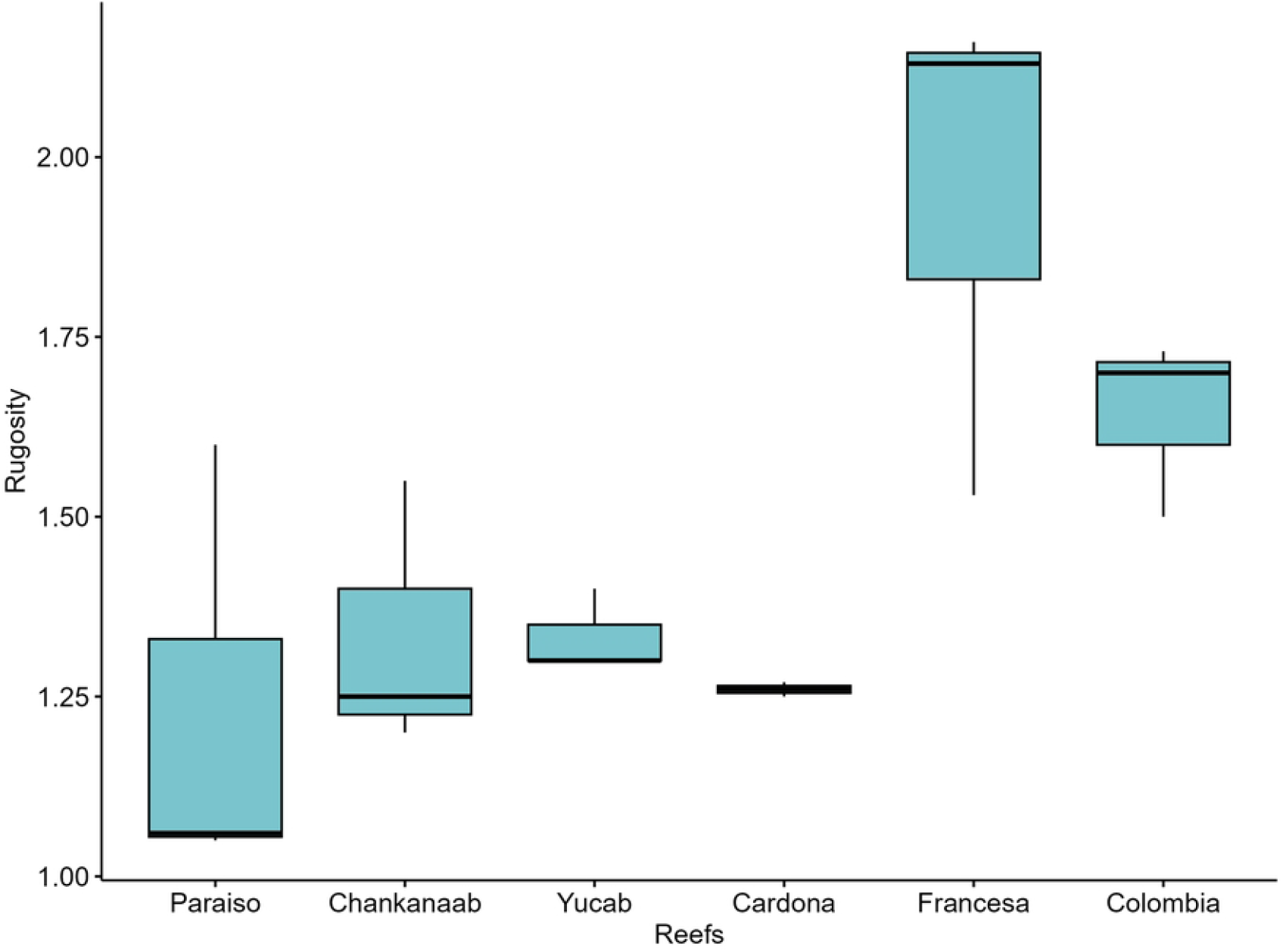
DGR values per reef. DGR was calculated only underneath of those areas where live corals were present. The ordering of reefs follows a north-to-south direction. Columns represent mean values, while bars indicate the standard error.

For total digital rugosity (DGR + DLR), all sites showed significant differences (p < 0.05, S3 File). The site Yucab exhibited the highest values, followed by Colombia, Francesa, Cardona, Chankanaab and Paraiso (Fig 4).

**Fig 4:**
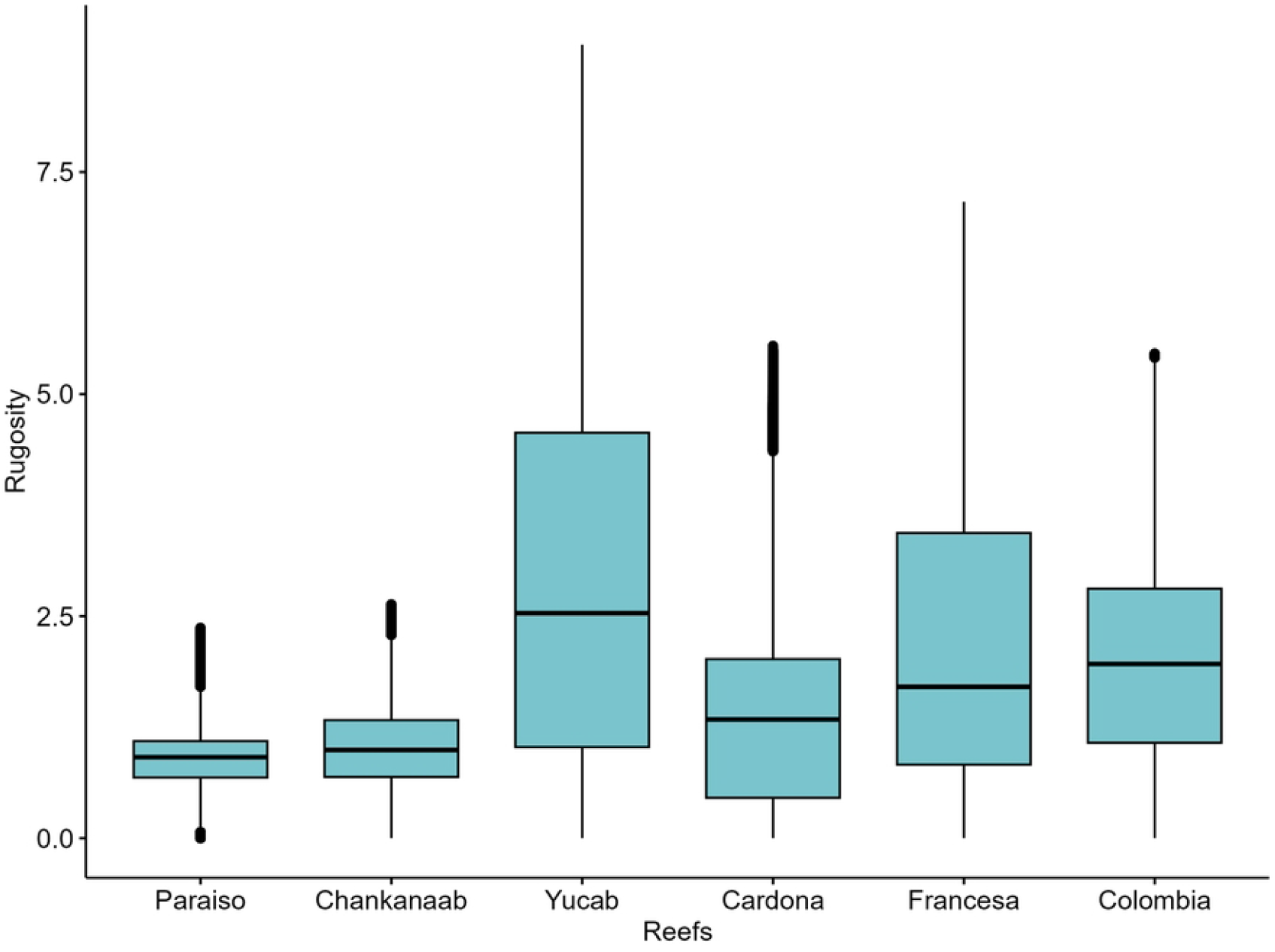
DLR values per reef, which is the rugosity contributed by live coral cover. The ordering of reefs follows a north-to-south direction. Columns represent mean values, while bars indicate the standard error.

When assessing DGR exclusively (Fig 5) in areas underneath live coral coverage, Yucab, Francesa, and Colombia reefs, had the highest values, whereas Paraíso showed the lowest rugosity, all sites were statistically different (p < 0.05, S4 File).

**Fig 5.**
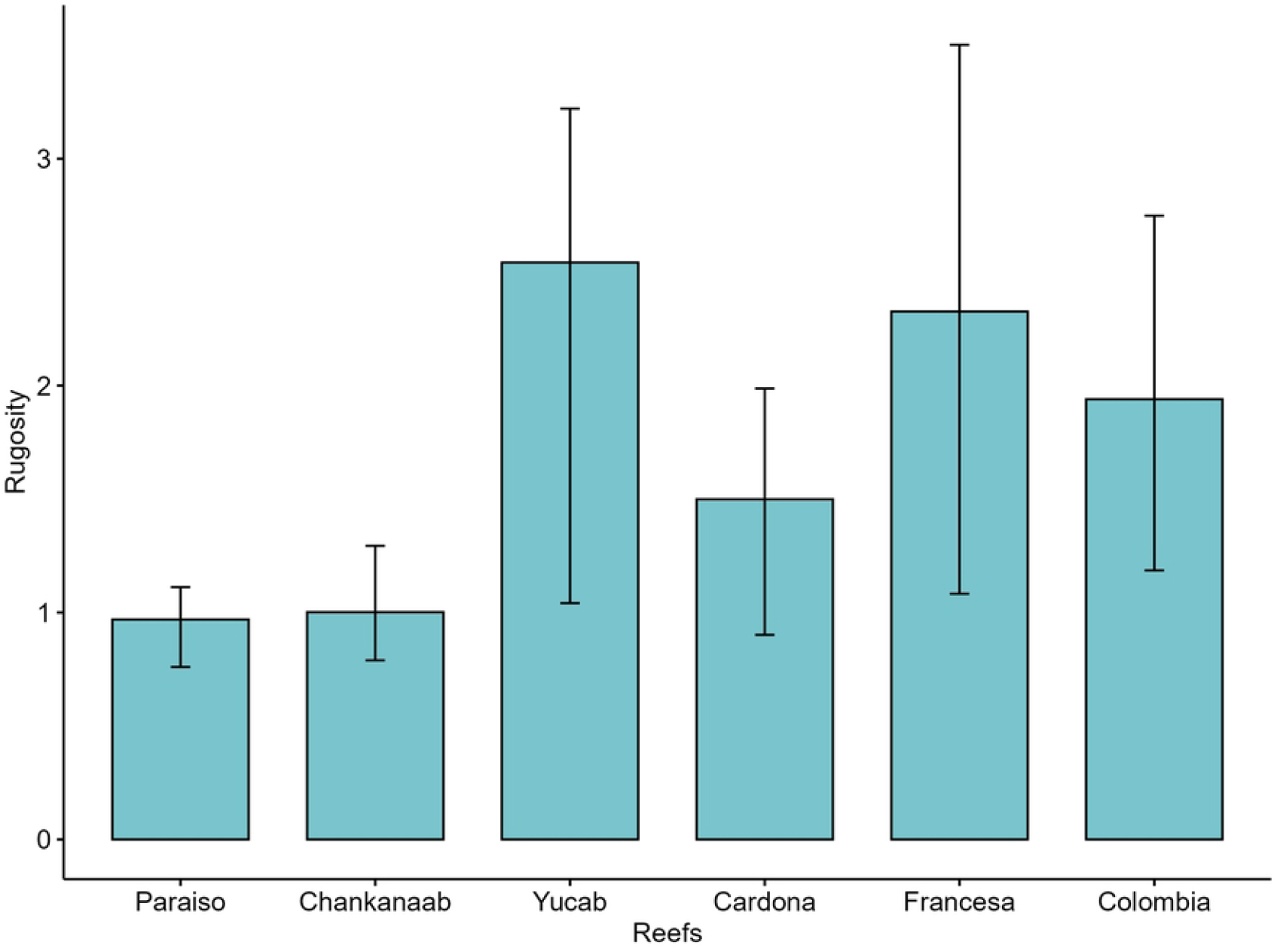
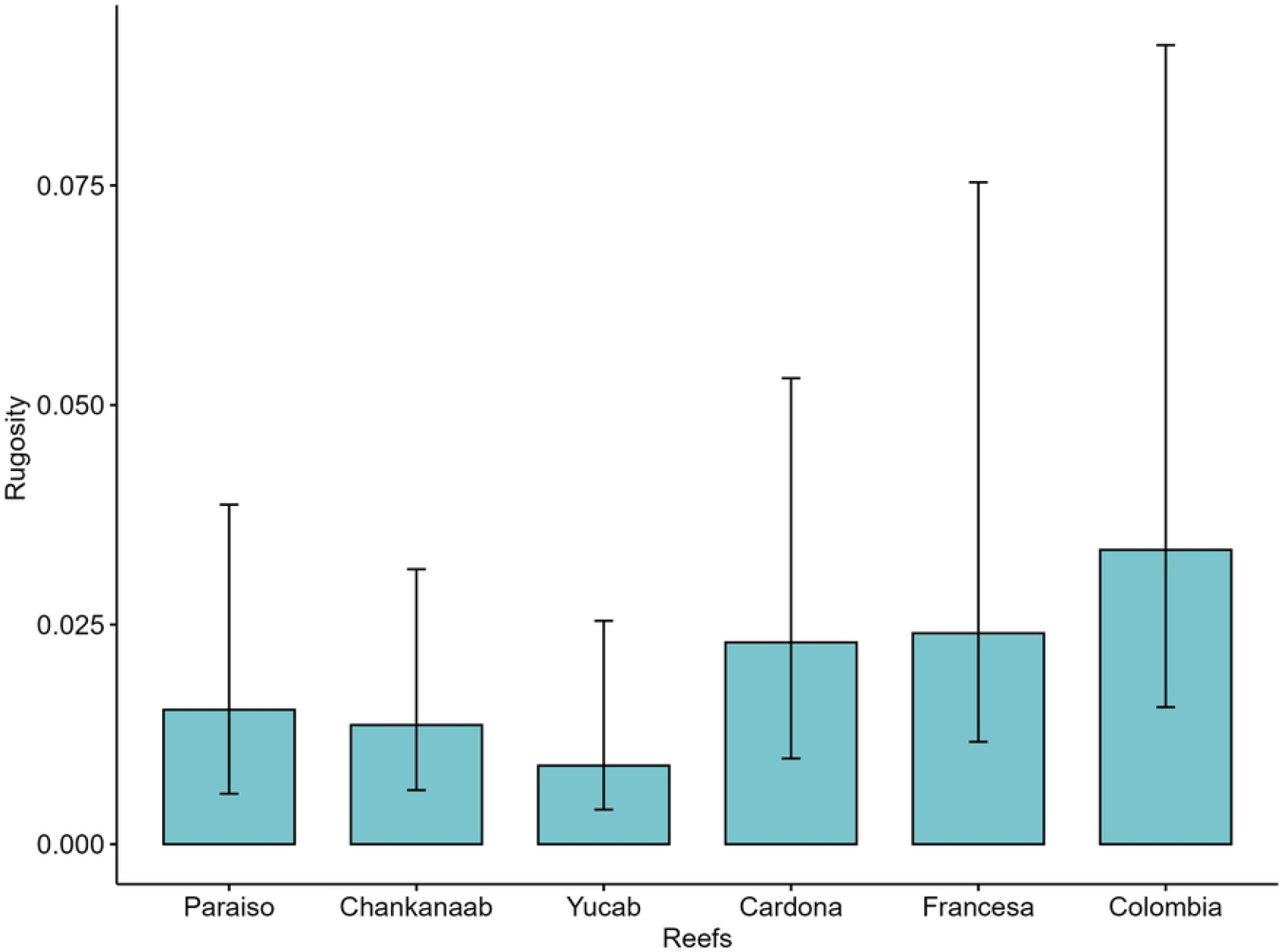
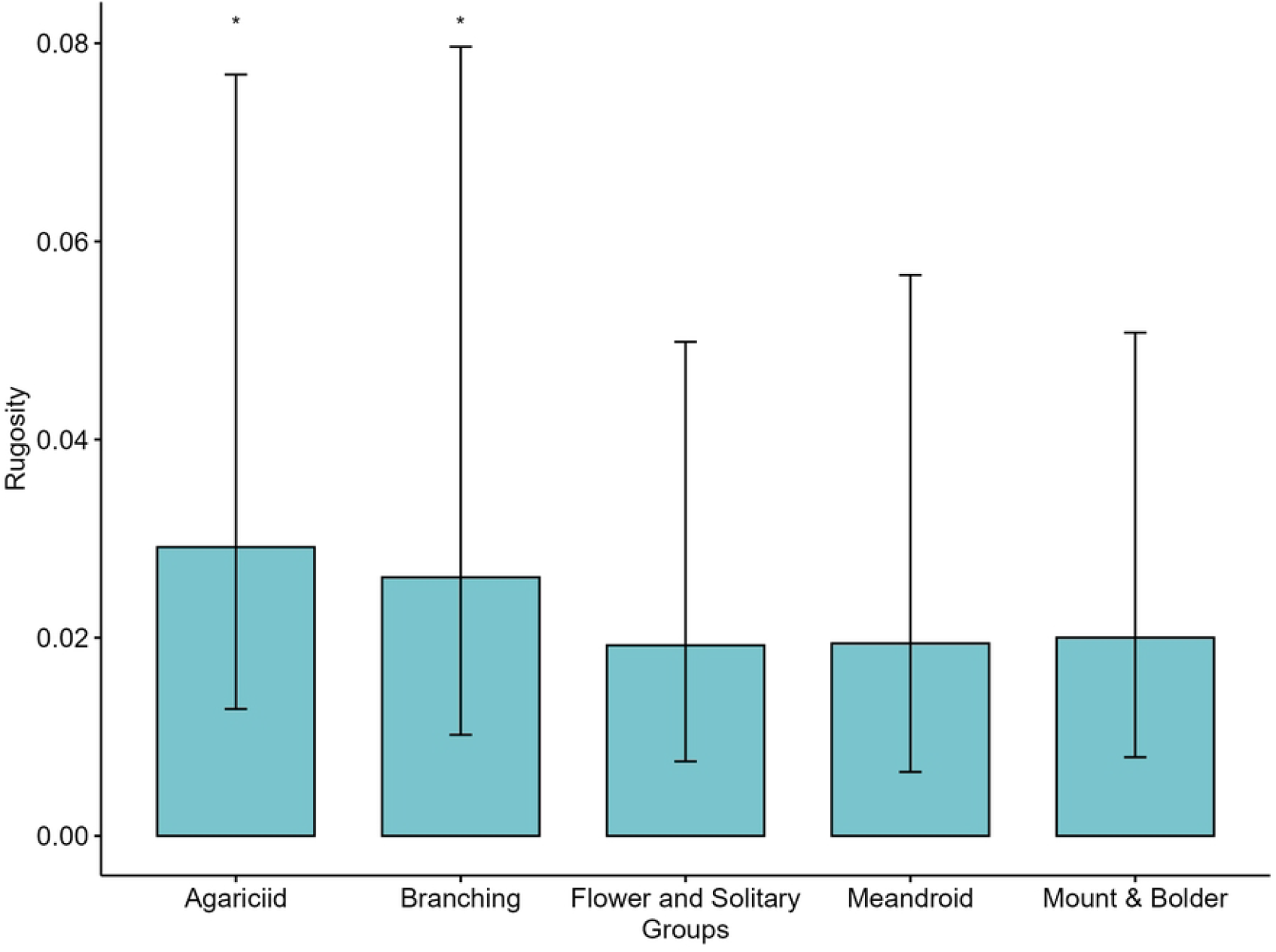
Contribution of coral groups to DLR in the CRNP. * Indicate statistically significant differences (p < 0.05), among coral groups.

For DLR (Fig 6), Colombia, Francesa, Cardona and Paraiso showed the highest values, whereas Yucab exhibited the lowest ones. All sites showed significant differences (p < 0.05, S5 File).

Colombia and Francesa reefs had the highest rugosity in both the DGR and DLR estimates. However, the results for Yucab were contrasting, as this reef presented the highest DGR, but the lowest DLR values.

In terms of contribution to DLR by coral morphological groups in the CRNP, the agariciid and branching groups are the highest contributors (p < 0.05). No significant differences were observed between Flower & solitary, Meandroid and, mount & bolder morphological groups (p > 0.05, S6 File).

## Discussion

Underwater digital photogrammetry improves the assessment of coral reefs features by allowing more accurate quantification of rugosity as a measurement of structural complexity, which is a key factor in understanding ecological processes and biodiversity patterns [22, 24, 44, 45].

We used Digital Elevation Models (DEMs) to accurately represent the reef rugosity across different scales. The statistical equivalence between the reconstructed and original DEMs confirmed the reliability of this approach. Discrepancies arised when comparing the rugosity assessments obtained from the *in situ* and digital methods.

The *in situ* approach, identified the Francesa, Colombia and Yucab reefs with the highest rugosity. In contrast, the total rugosity digital approach highlighted to Yucab, Colombia and Francesa reefs with the highest values. The chain method relies on physical measurements along specific transects, and its accuracy depends heavily on the number of replicates and the exact sites surveyed [46, 47]. A limited sampling can lead to under-representation of the spatial variability of the reef, potentially biasing the results if the surveyed areas are not representative of the entire reef [14, 48, 49]. This is not the case when using the approach based on UWP, as large areas are considered for the assessment.

Significant differences in rugosity exist among these reefs, driven by both DLR and DGR. This rugosity is shaped by benthic components, such as live coral coverage, as well as the calcareous matrix of the reef structure, collectively contributing to total rugosity. Digital underwater photogrammetry has allowed for a more precise quantification of rugosity [14, 24, 25], which is crucial for gaining a deeper understanding of the ecological processes within these habitats.

A clear distinction emerges in the interpretation of rugosity at both DLR and DGR levels, consistent with findings from studies such as Yanovski et al. (2017), who revealed that different spatial scales yield different patterns among sites. Ribas-Deulofeu et al. (2021) found that rugosity in coral reefs arises primarily from two sources, biotic components like hard corals and sponges, and coarse-scale topographic variations reflecting the geological context of the reefs.

In our study, the DLR was higher in Colombia, Francesa and Cardona reefs, and the lowest values were observed in Yucab. The DGR was higher in Yucab, Francesa and Colombia. The difference on the estimates for the Yucab reef, is likely related to the contribution from its calcareous matrix formed by the historical accretion of the scleractinian corals, reflecting the geological context [27, 47]. Yucab has the lowest coral cover (< 4%) compared to Colombia (c.a. 15%), Francesa (c.a. 4.55%) and Cardona (5.51 %). This suggests that Yucab was once a highly complex site with extensive live coral cover. The differences observed at various spatial scales show that Yucab, Colombia and Francesa, are the reefs with highest rugosity in the CRNP.

When focusing on DLR based solely on sites with coral coverage, the highest values were observed in Colombia, which also had the greatest coral cover [36]. DLR is primarily defined by corals with high coverage, with the dominant groups being agariciid corals followed by branching corals [36].

Agariciid corals, which currently contribute the most to the DLR, are considered “weedy” species [9, 50]. These species often quickly colonize disturbed areas and can dominate when competition for space is reduced, owing to the loss of other coral species [51]. Although they can increase coral cover in the short term, their dominance may indicate a degraded ecosystem [52].

Focusing solely on the total rugosity might lead to underestimation of the importance of coral coverage in maintaining rugosity over time. Our results, showed that the DGR (historical context) contributed the highest to the rugosity estimates. The chain method, accounts for the DLR and DGR, and not necessarily reflect the current context of coral coverage, as observed in Yucab reef.

Furthermore, while a reef may maintain a complex three-dimensional structure, low coral cover can negatively affect the density and diversity of other benthic organisms [53, 54]. This is a critical factor to consider when assessing reef health and functionality.

In the CRNP, the primary contributing groups to DLR are Agariciid and branching corals (p < 0.05), followed by mound & boulder corals and meandroid corals. Agariciid and branching corals, characterized by low relief, foliose, and ramified morphologies, are the dominant coral groups in the CRNP and have a limited contribution to the calcareous matrix [50, 52]. This indicates that key groups essential for supporting long-term structural complexity are scarce [15, 55]. Therefore, it is important to emphasize that relying solely on coral cover as a metric underestimates the responses of biological systems to both natural and anthropogenic pressures [56] which does not necessarily ensure the long-term physical functionality of reefs.

Our findings highlight the spatial variability in reef rugosity within the CRNP. The higher rugosity values in Yucab and Francesa suggest that these reefs may provide more complex habitats, support greater biodiversity, and offer more refuge for marine organisms [22, 57]. Conversely, the lower rugosity in Paraíso indicates a flatter reef structure, which may influence the types of species that inhabit these areas [2]. But assessing coral coverage and its contribution to DLR, as well as focusing on the principal groups that enhance DLR, can help to detect sites that may not support positive reef growth [3]. By mapping both DLR and DGR, assessing coral species composition, and identifying sites with high historical rugosity but low local biological rugosity, conservation efforts can be more effectively directed. Specifically, efforts can be focused on enhancing local rugosity through restoration, protecting existing structural foundations, and monitoring changes in the reef topography and biological composition over time.

## Conclusions

This study illustrates the effectiveness of a wavelet-based approach for separating high-resolution digital elevation models (DEMs) of coral reefs into detailed local rugosity (DLR) and general rugosity (DGR) components, enabling a comprehensive multiscale examination of reef structural complexity. By employing a wavelet filter on elevation profiles extracted from UWP derived DEMs, we were able to capture both fine-scale (DLR) and broad-scale (DGR) topographic features, providing a better understanding of reef rugosity than conventional methods.

Our results reveal that agariciid and branching corals are the main contributors to DLR in the CRNP. Although these “weedy” species can rapidly colonize disturbed areas, their prevalence may not support long-term structural complexity and could signal a degraded ecosystem. The lack of crucial coral groups necessary for maintaining long-term structural integrity underscores that coral cover alone is insufficient for evaluating reef health.

Reef areas exhibiting high DGR but low DLR, such as Yucab, may appear structurally complex but lack the living coral cover required to sustain diverse marine communities and to maintain vital ecosystem functions. Therefore, conservation initiatives should aim not only to preserve the physical structure of reefs, but also to improve local rugosity by increasing roughness through restoration and protection of coral species that contribute significantly to DLR.

A primary limitation of our methodology is the use of 2D format DEMs, which may not fully capture the intricacy of certain coral colonies, particularly those that grow in clumps. UWP provides information visible from the perspective of a camera, potentially overlooking complex three-dimensional structures. Future studies should investigate the application of this methodology to 3D models to represent the full complexity of coral reef structures.

## Acknowledgements

This research was funded by the PAPIIT grant (IN218219) from the National Autonomous University of Mexico supported this work. EB-F was supported by the CONACYT postgraduate scholarship.

## Supporting information

**S1 Fig**. Study Area, black dots indicate the locations of sampled reefs within the Cozumel Reefs National Park.

**S2a Fig**. Panel (a) illustrates DGR, highlighting the geological contributions to reef structure, while panel (b) depicts DLR, emphasizing the biological contributions from live coral communities,

**S2b Fig**. Panel (b) illustrates DGR, highlighting the geological contributions to reef structure, while panel (b) depicts DLR, emphasizing the biological contributions from live coral communities.

**S3 Fig**. Rugosity by reef using the chain method. The ordering of reefs follows a north-to-south direction. The black lines represent the median values of rugosity, highlighting variation among reefs.

**S4 Fig**. Total rugosity (DGR + DLR) per reef site. The ordering of reefs follows a north-to-south direction. This Fig integrates both the DGR representing the structural component shaped by long-term geological processes and DLR which reflects the contribution of living coral and benthic organisms.

**S5 Fig**. DGR considering only areas with coral coverage per reef site. The ordering of reefs follows a north-to-south direction. Bars represent mean values, while error bars indicate variability across measurements.

**S6 Fig**. DLR considering only areas with coral coverage per reef site. The ordering of reefs follows a north-to-south direction. Bars represent mean values, while error bars indicate variability across measurements.

**S7 Fig**.. Contribution of coral groups to DLR in the CRNP. Asterisks (*) indicate statistically significant differences (p < 0.05) among coral groups.

